# Direct shoot regeneration from cotyledon, leaf and root of *Citrus jambhiri* Lush

**DOI:** 10.1101/2020.09.30.320077

**Authors:** Priyanka Sharma, Bidhan Roy

## Abstract

*Citrus jambhiri* (Rough lemon) is popularly preferred for rootstock for cultivated species of *Citrus*. Tissue culture is an appreciable technique for mass-multiplication of plant propagules. In this communication direct regeneration of plantlets of *Citrus jambhiri* Lush. were obtained from cotyledons, roots and leaves. Most of the cotyledon (96%) enlarged on medium supplemented with 50 mg/L of casein hydrolysate. Few of those enlarged cotyledons responded to direct regeneration of shoots. Maximum shoot per responded cotyledon was 32. Conversely, the health of the plantlets were poor with semi-cylindrical leaves. Most of them dried on maintenance medium or on rooting medium ad died. Plantlets regenerated on medium supplemented with IAA in combination with IBA were healthy and they established on maintenance medium and rooted on rooting medium. Direct regeneration was also obtained from leaf on MS medium supplemented with 0.50 mg/L of dicamba. Our finding concluded that tissue culture tools may be used for direct regeneration of plantlets from different explants of *C. jambhiri* to obtained true-to-type plant propagules.

## Introduction

Citrus is one of the most popular edible fruit and it is being cultivated worldwide. Eastern Asia, particularly the south-east Asia is thought to be the primary centre of origin of *Citrus* species and in this region, many citrus species are still found in their wild state [1]. The north-eastern region of India is a part of the centre of origin and rich in diversity of citrus with wild and endangered species. It has wide range of uses, such as table purpose, processed products, culinary purpose as well as pharmaceutical products. A large number of different species are available under the genus-*C. jambhiri* (Rough lemon) is generally preferred for rootstock for lemons, oranges, mandarins, grape fruits and kinnows for its high vigour and wide adaptability well adaptation ability under problem soils and odd situations [2-4]. This species was found to have tolerant ability towards many biotic and abiotic stresses.

Direct regeneration of plantlets from somatic tissues hold extreme importance as it will produce true-to-type plantlets. There is ample changes of somaclonal variations in indirect regeneration of plantlets through callus induction. Thus, direct regeneration of plantlets from explant may play important role in mass-multiplication of true-to-type plant propagules. There are large number of research works on direct regeneration from nodal segment and shoot tips of *Citrus* species [1, 5-7]. However, limited research findings are available on direct regeneration from cotyledon, leaf and root of *Citrus* spp. Contrariwise, plenty successful research findings are available on indirect regeneration through callus initiation and plantlet regeneration [3, 7-12]. Hence, in this endeavour effort was taken to standardize protocol for direct regeneration of plantlets from cotyledon, root and leaf of *Citrus jambhiri* Lush.

## Materials and methods

### Plant materials

Collected mature fruits of *C. jambhiri* Lush. were cut with sharp knife and seeds were extracted manually. Seeds were surface sterilized with 0.1% HgCl_2_ for 10 minutes followed by 3-5 times washing with sterilized distilled water.

### Medium preparation

Medium was prepared using all of the individual chemical compounds listed by Murashige and Skoog [13]. Individual stock solutions were prepared and stored in separate bottles for ready use during the preparation of culture media. MS medium was prepared with 3% sucrose and 0.8% agar. The pH of medium was adjusted to 5.8. The media were autoclaved at 121 °C under 104 kPa for 15 minutes for sterilization.

### Inoculation for direct regeneration from cotyledons

Surface sterilized seeds were inoculated on MS medium fortified with different concentrations of casein hydrolysate and different concentrations and combinations of growth regulators as listed in Table 1. Cultures were then incubated in culture room at 25±2 °C with 16/8 h light and dark phases for six weeks.

**Table 1.** Effect of different concentrations and combination of growth regulators on direct regeneration from cotyledon of *C. jambhiri* on casein-hydrolysate supplemented MS media.

| Treatments | No. of seeds produced enlarged cotyledons | Average weight of enlarged cotyledons (g) | No. of cotyledon produced multiple shoots | Sl. No. of cotyledon produced multiple shoots | No. of multiple shoots/ cotyledon |  |
| --- | --- | --- | --- | --- | --- | --- |
|  |  |  |  |  | Individual cotyledon | Mean |
| T1 | 48 (96%) | 1.628 a | 3.0 | 1 | 32.0 a | 15.0 a |
|  |  |  |  | 2 | 12.0 b |  |
|  |  |  |  | 3 | 1.0 e |  |
| T2 | 42 (84%) | 1.148 b | 0.0 | - | 0.0 f | 0.0 c |
| T3 | 39 (78%) | 1.050 b | 0.0 | - | 0.0 f | 0.0 c |
| T4 | 28 (56%) | 0.517 d | 0.0 | - | 0.0 f | 0.0 c |
| T5 | 32 (64%) | 0.553 d | 0.0 | - | 0.0 f | 0.0 c |
| T6 | 37 (74%) | 0.815 c | 4.0 | 1 | 3.0 d | 3.0 b |
|  |  |  |  | 2 | 1.0 e |  |
|  |  |  |  | 3 | 7.0 c |  |
|  |  |  |  | 4 | 1.0 e |  |
| T7 | 13 (26%) | 0.682 e | 0.0 | - | 0.0 f | 0.00 |
| Total | 239 | - | 7.0 | - | - | - |
| Range | 13-48 (96-26%) | 0.517-1.628 | 0.0-4.0 | - | 1.0-32.0 | 3.0-15.0 |
| Mean | 34.14 (68.29%) | 0.76 | 3.50 | - | 8.14 | 9.0 |

**T1:** Casein-Hydrolysate @ 50 mg/L; **T2:** Casein-Hydrolysate @ 100 mg/L; **T3:** Casein-Hydrolysate @ 200 mg/L; **T4:** IAA @ 1.0 mg/L; **T5:** IBA @ 1.0 mg/L; **T6:** IAA @ 1.0 mg/L + IBA @ 1.0 mg/L; **T7:** BAP @ 1.0 mg/L.
\*Values bearing same letter in the column are not significantly different at $p = 0.05$ of LSD

### Inoculation for direct regeneration from root

Similarly, the surface sterilized seeds were inoculated on MS medium added with different concentrations and combinations of IAA and IBA as listed in Table 2. Cultures were then incubated in culture room at 25±2 °C with 16/8 h light and dark phases for six weeks.

**Table 2.** Effect of growth regulator on direct shoot regeneration from root of *Citrus jambhiri*.

| Treatment |  | No. of seed<br>inoculated | No. of seedling<br>responded to direct<br>regeneration from root | % of<br>response | Average No. of<br>shoots per<br>inoculated shoot |
| --- | --- | --- | --- | --- | --- |
| IAA | IBA |  |  |  |  |
| 0.50 | 0.50 | 60 | 0.0 | 0.00 | 0.00 |
| 1.00 | 0.50 | 60 | 0.0 | 0.00 | 0.00 |
| 0.50 | 1.00 | 60 | 0.0 | 0.00 | 0.00 |
| 1.00 | 1.00 | 60 | 2.0 | 3.33 | 4.50 |
| 1.00 | 1.50 | 60 | 0.0 | 0.00 | 0.00 |
| 1.00 | 2.00 | 60 | 0.0 | 0.00 | 0.00 |
| 1.50 | 1.00 | 60 | 0.0 | 0.00 | 0.00 |
| 2.00 | 1.00 | 60 | 0.0 | 0.00 | 0.00 |

### Inoculation for direct regeneration from leaf

Surface sterilized seeds inoculated on MS basal medium for germination and establishment of seedling. Seeds in culture bottles were kept in culture room at 25±2 °C with 16/8 h light and dark phases. Six week old *in vitro* established seedling were used as source material for leaf explant. Leaves were excised from seedlings under laminar air flow cabinet. Leaves were injured with the help of the back side of the scalpel. The leaves were inoculated on MS medium fortified with different concentrations and combination of growth regulators as listed in Table 3. The cultures were again kept in culture room at 25±2 °C with 16/8 h light and dark phases.

**Table 3.** Effect of different concentrations and combinations of growth regulators on direct shoot regeneration from leaf of *C. jambhiri*.

| Treatments | Callus induction (%) | Regeneration on callus induction medium |
| --- | --- | --- |
| 2,4-D 1.0 mg/L | 78.57 | - |
| 2,4-D 2.0 mg/L | 64.51 | - |
| 2,4-D 1.0 mg/L + NAA @ 0.50 mg/L | 100.00 | - |
| 2,4-D 2.0 mg/L + NAA 0.50 mg/L; | 100.00 | - |
| Picloram 0.50 mg/L | 88.88 | - |
| Picloram 1.0 mg/L | 90.91 | - |
| Dicamba 0.50 mg/L | 84.00 | 9.75% |
| Dicamba 1.0 mg/L | 76.92 | - |
| TDZ 0.25 mg/L | 96.87 | - |
| TDZ 0.50 mg/L | 100.00 | - |

## Results

### Direct Regeneration from Cotyledon

The experiment was set to study the efficiency of direct shoots regeneration from cotyledons of germinating seeds of *C. jambhiri*. During the process of germination of seeds, the cotyledon enlarges. It varied from 96% to 26% (Table 1). High response towards enlargement of cotyledon was observed in medium fortified with casein hydrolysate. Medium added with 50 mg/L of casein hydrolysate produced maximum number of enlarged cotyledons (96%) followed by 100 mg/L of casein hydrolysate (84%) and 200 mg/L of casein hydrolysate (78%). Average weight of enlarged cotyledons varied from 0.517 to 1.628 g/cotyledon with a mean of 0.76 g/cotyledon (Table 1). Largest cotyledons by weight were achieved on the synthetic medium supplemented with 50 mg/L of casein hydrolysate (1.628 g/cotyledon) followed by 100 mg/L of casein hydrolysate (1.148 g/cotyledon) and 200 mg/L of casein hydrolysate (1.050 g/cotyledon).

Direct regeneration from enlarged cotyledon was observed when MS medium supplemented with 50 mg/L of casein-hydrolysate (Fig. 1A,B&C) and IAA of 1.0 mg/L + 1.0 mg/L of IBA (Fig. 2A,B,C&D). However, the direct regeneration from cotyledon was random, only seven cotyledons produced direct shoots (Table 1), three on medium supplemented with 50 mg/L of casein-hydrolysate and four on medium supplemented with 1.0 mg/L of IAA + 1.0 mg/L of IBA. The seeds that were inoculated on the medium fortified with 50 mg/L of casein hydrolysate, after three weeks of inoculation, numerous globular growth was observed on the enlarged cotyledon (Fig. 1A&B). Few of those globular growth regenerated into plantlets without roots. Leaves of regenerated plantlets were not normal (Fig. 1A&B), it was narrow and cylindrical (Fig. 1D).

**Fig. 1.**
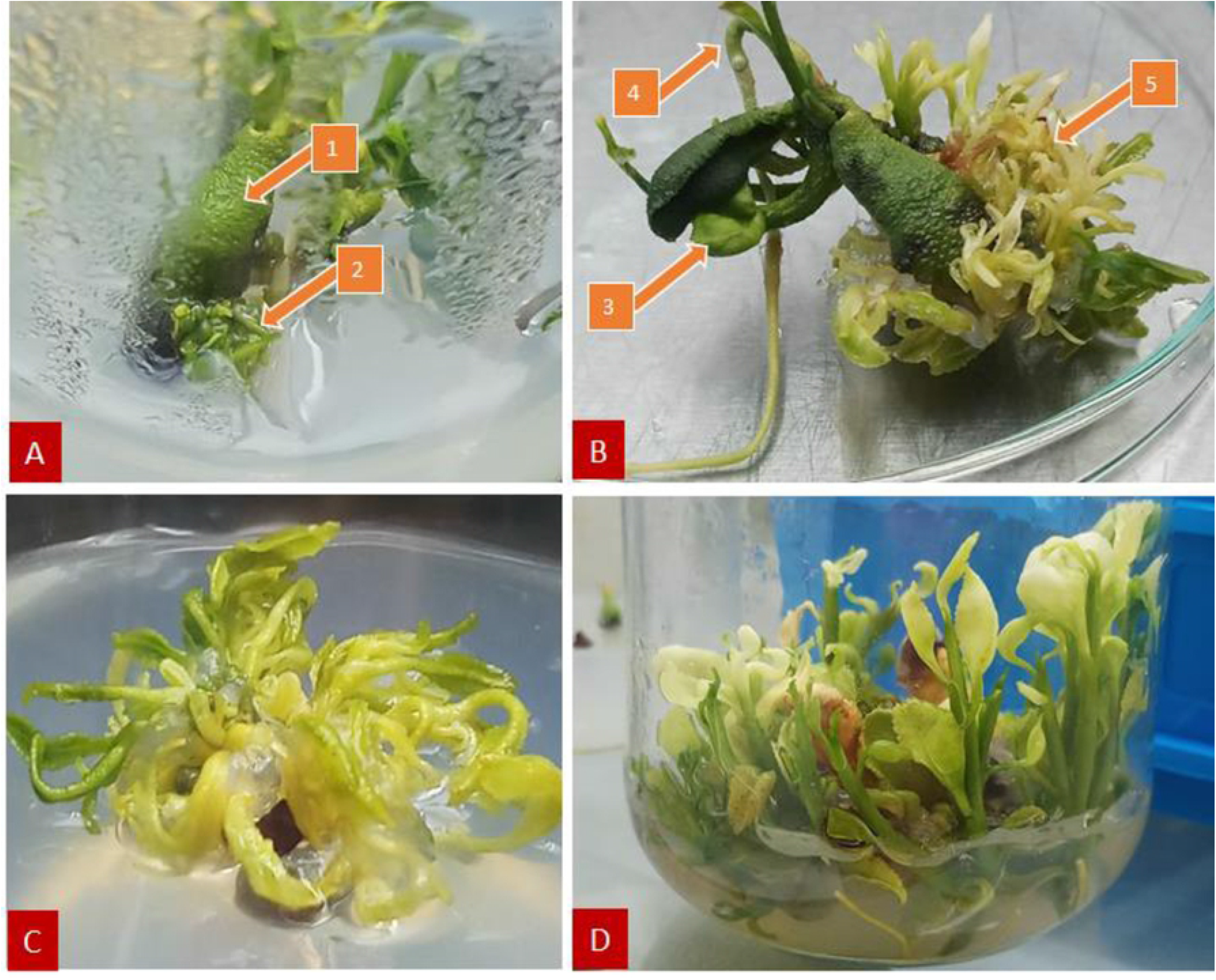
Direct regeneration from cotyledons of *Citrus jambhiri* when inoculated on MS supplemented with 50 mg/L of casein-hydrolysate. **A)** Initiation of multiple buds on cotyledon: **1]** Enlarge cotyledon, **2]** Multiple sprouted buds; **B)** Grownup of multiple plantlets on cotyledon: **3]** Plumule developed from the seed, **4]** Radicle (root) developed from the seed, **5]** Grownup of multiple plantlets; **C)** Grownup of multiple plantlets were separated from cotyledon and again culture on MS basal medium; **D)** Multiple plantlets regeneration from cotyledonary axil of germinating seeds when inoculated on MS medium supplemented with 50 mg/L of casein hydrolysate.

**Fig. 2.**
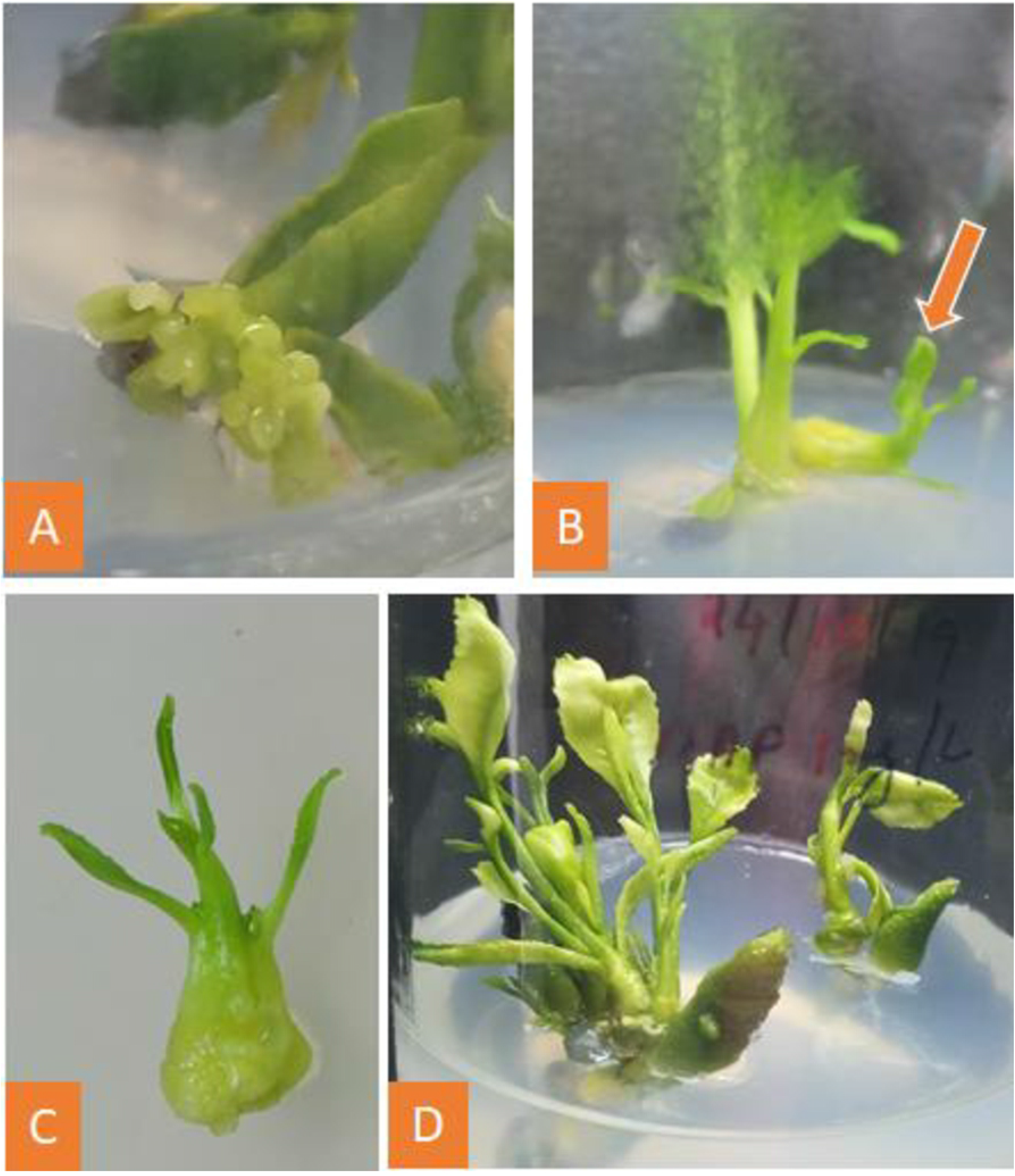
Direct regeneration from cotyledons of *Citrus jambhiri*. **A)** Direct embryogeneis on MS medium fortified with 1 mg/L of IAA and 1 mg/L IBA; **B)** Shoot regeneration on MS medium fortified with 1 mg/L of IAA and 1 mg/L IBA; **C)** Magnified portion of regenerating shoot from cotyledon; **D)** Multiple shoot regeneration on IAA and 1 mg/L IBA.

In contrast, the cotyledon enlarged on IAA @ 1.0 mg/L + IBA @ 1.0 mg/L supplemented medium produced prominent globular callus like structure with light green pigmentation (Fig. 2A). Gradually those globular structure converted into plantlets (Fig. 2B,C&D). The regenerated plantlets on IAA @ 1.0 mg/L + IBA @ 1.0 mg/L fortified medium were normal as that of plantlets regenerated from the cotyledonary axis. Highest number of shoots (32 shoots/cotyledon; Fig. 1B; Table 1) was recorded on 50 mg/L of casein-hydrolysate followed by 12 shoots/cotyledon on the medium with same growth regulators (Fig. 2D; Table 1).

### Direct regeneration from root

Direct regeneration from the root of germinating seeds were observed in *Citrus jambhiri* when seeds were inoculated on MS added with 1.0 mg/L of IAA and 1.0 mg/L IBA (Fig. 3 A,B&C). This event was unintended and random. Germination of seeds was usual, producing radicle and plumule simultaneously (Fig. 3A[1]). The tap root of germinating seed grew about 2.50 cm without touching the medium (Fig. 3A[2]). Direct regeneration of shoot and root were observed when it touched medium (Fig. 3B&C[3]). Only two root showed direct regeneration (Table 2). The direct *in vitro* regenerated shoots were hardened and finally transferred to the field as per the guidelines of [7].

**Fig. 3.**
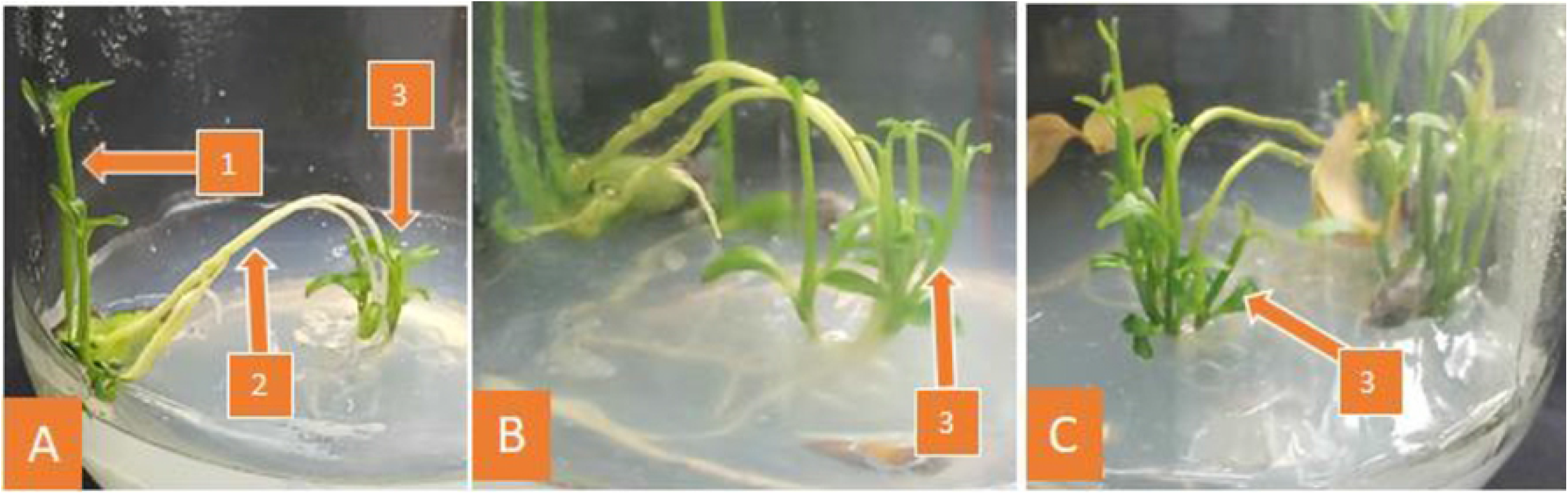
Direct regeneration from roots of *Citrus jambhiri*. **A**,**B&C)** Direct regenerated plantlets: **1]** Plumules from the germinated seeds, **2]** Tap root of germinating seed grew about 2.50 cm without touching the medium, **3]** Shoot regenerated directly from the root.

### Direct regeneration from leaf

A protocol was developed for direct plantlet regeneration from *in vitro* regenerated leaf explants of *C. jambhiri*. Leaves excised from axenic shoot cultures were used to induce organogenesis on MS medium added with different combinations and concentrations of growth regulators (Table 3). Medium added with 0.50 mg/L of dicamba showed very small callus on the leaf. Gradually the leaf dried up and the callus showed regeneration on the callus induction medium (Fig. 4B). Only 9.75% of the leaf callus on medium fortified with 50 mg/L of dicamba showed this type of regeneration (Table 3).

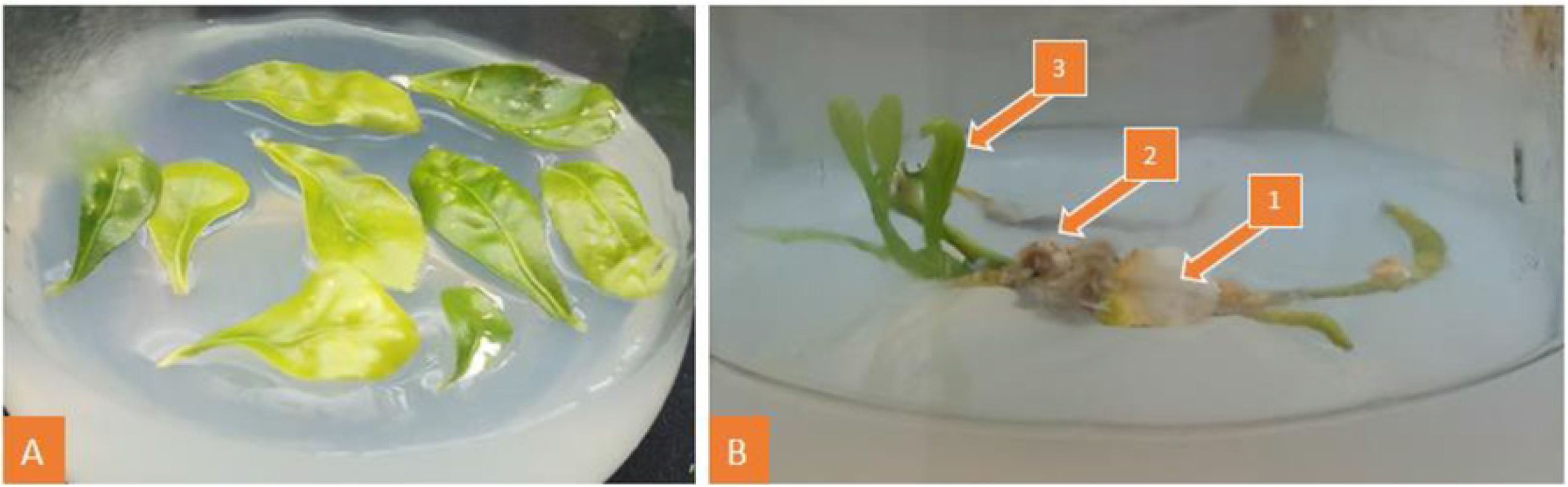

## Discussion

Direct regeneration of plantlets is commonly practiced for *in vitro* mas-multiplication of citrus because it ensures maximum genetic uniformity of the resulting plants [1, 14]. Direct regeneration from cotyledon of *C. jambhiri* is the novel finding of this endeavour. There are lot of research findings of indirect regeneration (through callus induction and shoots regeneration) form cotyledon derived callus of citrus species [3, 8, 10-12]. Contrariwise, as per the search for the literature in ‘Google Search’, there is no finding on direct regeneration of shoots from cotyledon of citrus species. Ample references are also available on direct shoot multiplication from nodal segments [1, 6, 7, 9, 15, 16], shoot tips [6, 7, 16, 17], cotyledonary node [18, 19], axillary buds [20] and meristem culture [21] of different citrus species.

Direct regeneration from cotyledon was obtained when the MS medium was supplemented with 50 mg/L of casein hydrolysate and 1.0 mg/L of IAA in combination with 1.0 mg/L of IBA. Large number of plantlets regenerated from cotyledon when the MS medium was supplemented with 50 mg/L of casein hydrolysate, but the health of the plantlets were weak with mostly semi-cylindrical leaves. Casein hydrolysate overcomes the shortage of glutamine when there is insufficient phosphorus for adequate biosynthesis however several investigators have concluded that casein hydrolysate itself is more effective for plant culture than the addition of the major amino acids. This has led to assumption that casein hydrolysates might contain some unknown growth promoting factor [22].

Most of the plantlets obtained on casein hydrolysate added medium dried during subsequent course of the culture. Only few of the plantlets survive and rooted on the rooting medium. Yet, the plantlets regenerated on 1.0 mg/L of IAA in combination with 1.0 mg/L of IBA added medium were normal. Plantlets were rooted and planted in field after hardening.

Direct plantlets regeneration from root also very scanty. Bhat et al. [23] recorded *de novo* shoot bud initiation in basal medium at a low frequency during three years of continuous culture of roots of *Citrus aurantifolia* (Christm.) Swing. There are ample research findings on indirect regeneration (through callus induction and shoots regeneration) from root derived callus of citrus species [2, 9, 24]. However, research findings on direct regeneration from root of *C. jambhiri* is not available. Thus, our research finding on direct regeneration of plantlet from root is innovative.

Direct plantlet regeneration from leaf segments represent a promising tool for mass-multiplication of citrus keeping the genetic fidelity intact. To date, direct organogenesis from leaf explants of *C. jambhiri* is not available. Explants on a medium with a high concentration of cytokinin-to-auxin ratio, they will develop buds/shoots [25-27]. Kasprzyk-Pawelec et al. [28] also reported this type of direct organogenesis from leaf explant of *C. limon* L. Burm cv. ‘Primofiore’ when the leaf explants were cultured on MS medium supplemented with 3.5 m/L of BAP. Finding of Hu et al. [29] correspondingly suggested that cytokinin was the primary factor for shoot organogenesis in citrus. However, some reports are available on direct regeneration from leaf of plant species other than citrus. Varutharaju et al. [30] standardized an efficient protocol for direct plantlet regeneration has for the medicinal plant *Aerva lanata* (L.) Juss. ex Schult. An efficient propagation and regeneration system through direct plantlet organogenesis from leaf explant was established in *Lysionotus serratus* by Li et al. [31]. They found that high concentration of 6-benzyladenine (BA) or thidiazuron (TDZ) was effective for direct organogenesis. Tilkat et al. [32] developed a protocol for direct plantlet regeneration from leaf explants of male *Pistacia vera* L. cv. ‘Atli’. Leaves excised from axenic shoot cultures of pistachio were used to induce organogenesis on MS medium with Gamborg vitamins added with different combinations and different concentrations of BAP and IAA. Bobatk et al. [33] obtained direct shoot organogenesis of *Drosera rotundifolia* L. on MS basal medium or MS medium supplemented with 10-8 M NAA. Liquid culture medium significantly increased regeneration capacity of leaf tissue. Their findings on direct shoot organogenesis was supported by histological and scanning electron microscopy investigations and it was found that direct plant regeneration was without intermediate callus formation.

## Conclusion

Direct regeneration were achieved from cotyledons, roots and leaves. The regeneration obtained from leaves was through callus induction, but plantlets induction taken place on callus induction medium. The plantlets obtained from cotyledons on MS medium supplemented with 0.5 mg/L of casein hydrolysate were poor in health and most of the plantlets died on maintenance medium or on rooting medium. Only few of them survived. The cotyledonary plantlets obtained on IAA and IBA supplemented medium were normal in health all the regenerated plantlets survived and rooted on rooting medium. Plantlets regeneration from root was random. Only 9.75% *in vitro* growing seedlings responded to direct plantlets regeneration. The plantlets regenerated from roots were in good health and they all survived and rooted on rooting medium. Our findings established that the tissue culture tool may be used for direct regeneration to obtain true-to-type plant propagules.

